# ReaxFF-guided Optimisation of VIRIP-based HIV-1 Entry Inhibitors

**DOI:** 10.1101/2025.02.11.637598

**Authors:** Fabian Zech, Christoph Jung, Armando Alexei Rodríguez-Alfonso, Janet Köhler, Ludger Ständker, Gilbert Weidinger, Timo Jacob, Frank Kirchhoff

**Affiliations:** Institute of Molecular Virology, Ulm University Medical Center, 89081 Ulm, Germany; Institute of Electrochemistry, Ulm University, 89081 Ulm, Germany; Helmholtz-Institute Ulm (HIU) Electrochemical Energy Storage, 89081 Ulm, Germany; Karlsruhe Institute of Technology (KIT), 76021 Karlsruhe, Germany; Core Facility Functional Peptidomics, Ulm University Medical Center, 89081 Ulm, Germany; Core Unit Mass Spectrometry and Proteomics, Ulm University Medical Center, 89081 Ulm, Germany; Institute of Biochemistry and Molecular Biology, Ulm University, Ulm 89081, Germany

**Keywords:** HIV-1, VIRIP, peptide fusion inhibitor, viral fusion, peptide therapeutics

## Abstract

Peptides hold great promise for safe and effective treatment of viral infections. However, their use is often constrained by limited efficacy and high production costs, especially for long or complex peptide chains. Here, we used ReaxFF molecular dynamics (MD) simulations to optimize the size and activity of VIRIP (Virus Inhibitory Peptide), a naturally occurring 20-residue fragment of α1-antitrypsin that binds the HIV-1 GP41 fusion peptide (FP), thereby blocking viral fusion and entry into host cells. Specifically, we used the NMR structure of the complex between an optimized VIRIP derivative (VIR-165) and the HIV-1 gp41 FP for ReaxFF-guided *in silico* analysis, evaluating the contribution of each amino acid in the interaction of the inhibitor with its viral target. This approach allowed us to reduce the size of the HIV-1 FP inhibitor from 20 to 10 amino acids (2.28 to 1.11 kDa). HIV-1 infection assays showed that the size-optimized VIRIP derivative (soVIRIP) retains its broad-spectrum anti-HIV-1 capability and is non-toxic in the vertebrate zebrafish model. Compared to the original VIRIP, soVIRIP displayed more than 100-fold higher antiviral activity (IC_50_ of ∼120 nM). Thus, it is more potent than a dimeric 20-residue VIRIP derivative (VIR-576) that was proven safe and effective in a phase I/II clinical trial. Our results show that ReaxFF-based MD simulations represent a suitable approach for the optimization of therapeutic peptides.

## Introduction

The development of peptide drugs, typically composed of short amino acid sequences between 0.5 and 5 kDa, made great progress in recent years and has enormous therapeutic potential.^1,2^ Since the discovery of insulin in 1921, more than 80 peptide drugs have been clinically approved, revolutionizing the treatment of conditions like diabetes.^3^ Early peptide drugs, such as insulin and adrenocorticotrophic hormone, were derived from natural sources.^3^ However, advances in protein purification and synthesis during the 20th century facilitated the development and application of synthetic peptides.^2^ Recent technological progress in structural biology and recombinant technologies has further accelerated the development of peptide therapeutics.^4,5^ These drugs span a wide range of therapeutic applications from metabolic and cardiovascular diseases to oncology and antimicrobial treatments.^2,6–8^ More than 170 peptides are currently in clinical development and many more in preclinical stages,^2,3^ highlighting the increasing significance and ongoing innovation in this field.

Peptide drugs are also a promising class of therapeutics against viral pathogens.^9–13^ The entry process of enveloped viruses presents an excellent target for antiviral therapy because it involves several steps that can be interrupted by different classes of inhibitors.^14^ In addition, preventing viral entry reduces the risk of harmful inflammatory responses and cell death.^15,16^ Many viral pathogens enter host cells through similar mechanisms involving receptor binding, fusion peptide exposure, six-helix bundle formation, and membrane penetration.^17,18^ For example, SARS-CoV-2 initially binds to the ACE2 receptor, while HIV-1 first binds to the cluster of differentiation 4 (CD4) receptor and subsequently engages the co-receptors CCR5 or CXCR4. These interactions trigger conformational changes in the respective envelope glycoproteins resulting, in the exposure of the fusion peptide (FP). This hydrophobic peptide then inserts into the host cell membrane, stably anchoring the virion to the target cell.^19,20^ Finally, helical N- and C-terminal heptad-repeat sequences form a six-helix bundle, which pulls the viral and host membranes together to mediate viral entry. The first clinically approved peptidic HIV-1 entry inhibitor, Enfuvirtide (Fuzeon, T20), prevents six-helix bundle formation.^21,22^ More recently, highly potent peptides acting by the same mechanism to inhibit SARS-CoV-2 and other coronaviruses have been reported^23,24^ and are currently evaluated in clinical trials.^25^

The VIRus Inhibitory Peptide (VIRIP) targets the step just before six-helix bundle formation, *i*.*e*. insertion of the HIV-1 gp41 FP into the cell membrane.^26^ VIRIP was initially discovered by screening of a human hemofiltrate peptide library and represents a naturally-occurring 20 amino acid residue fragment of α1-antitrypsin, the most abundant circulating serine protease inhibitor.^26^ VIRIP specifically binds to the highly conserved FP region of the HIV-1 transmembrane glycoprotein 41 (gp41), thereby blocking its penetration into the host cell membrane and preventing viral entry.^26^ This mechanism came as a surprise because it was previously thought that viral FPs shoot into the membrane like harpoons and that this step is too fast to be targeted.^27^ However, after the identification of the HIV-1 FP as a site of vulnerability, it has been shown that vaccination with the FP induces broadly neutralizing antibodies.^28^ Structure-function analyses allowed to greatly enhance the antiviral activity of VIRIP.^26,29,30^ In addition, VIRIP-based inhibitors proved to be active against all HIV-1 variants and did not show cross-resistance with other antiretroviral drugs. Resistance to improved VIRIP derivatives has a very high genetic barrier and comes at enormous costs for viral replication fitness.^31,32^ Most notably, monotherapy with an optimized analogue (VIR-576) was safe and effective in a phase I/II clinical trial.^33^ However, the necessity of intravenous injection of high doses of VIR-576, together with the availability of various highly active and orally available drugs against HIV-1, made VIRIP-based inhibitors unattractive for broad therapeutic use in humans. To overcome some of these drawbacks and to pave the way for optimization of other peptide drugs with defined targets, we applied a ReaxFF-guided strategy to further optimize the size and activity of VIRIP.

## Results and discussion

VIRIP binds to the HIV-1 gp41 FP, thereby preventing the anchoring of the viral particle to the cellular membrane and consequently its entry into the target cells (Fig. 1A, upper panel). Based on the NMR structure of VIR-165, an optimized VIRIP derivative, in complex with the HIV-1 gp41 FP (PDB 2JNR),^26^ we utilized the rotamer function of UCSF Chimera to predict the conformation of VIR-148 (Fig. 1A, lower panel). The latter differs from VIR-165 by a single amino acid change (F12V) and was selected because it has an IC_50_ of 180 nM, making it the most active VIRIP derivative identified in comprehensive structure-activity-relationship (SAR) studies.^26^ ReaxFF-based molecular modelling revealed that binding of the HIV-1 FP is mainly mediated by amino acids near the N and C termini of VIR-148 (Fig. 1B). This mode of binding is supported by alanine exchange SAR studies of the original VIRIP peptide.^26^ It has been previously shown that introduction of cysteine residues substantially enhances the antiviral activity of VIRIP derivatives.^26^ Notably, the NMR structure reveals that establishment of a cysteine bridge forces VIR-148 to form a loop structure that likely stabilizes the conformation allowing effective binding of the HIV-1 gp41 FP (Fig. 1A, lower panel).

**Figure 1.**
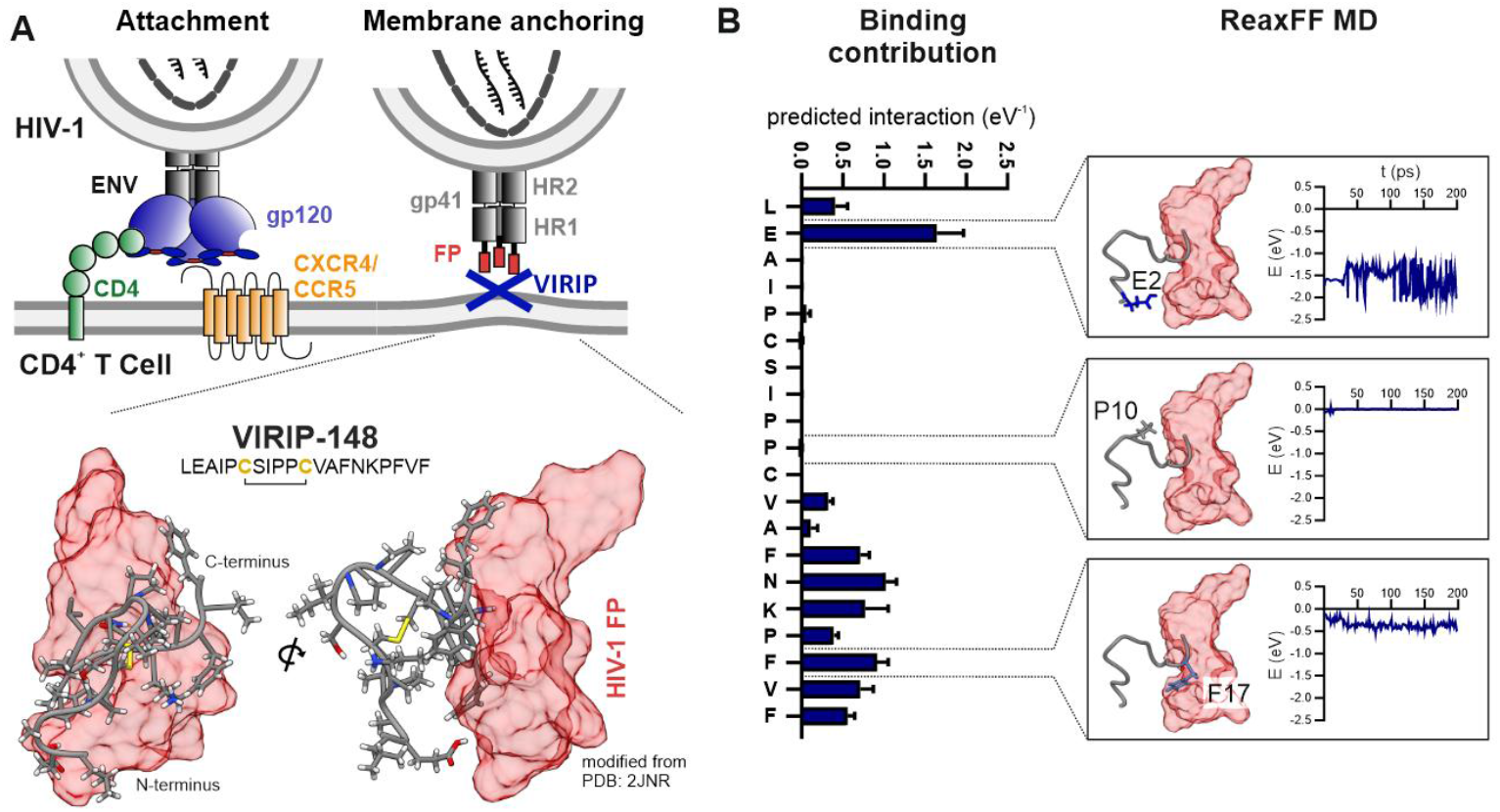
C- and N-terminal interactions mediate binding of VIR-148 to the HIV-1 gp41 FP. (A) Schematic presentation of HIV-1 attachment and membrane anchoring (upper panel). HIV-1 binds to CD4 and a co-receptor (CXCR4 or CCR5) on CD4+ T cells. The viral gp41 FP (red) is blocked by VIRIP. The lower panel shows VIR-148 (modified from PDB 2JNR^26^) bound to the HIV-1 gp41 FP. (B) Predicted contribution of each amino acid in VIR-148 to binding of the gp41 FP (left panel). ReaxFF MD simulations for selected Alanine mutations: E2A, P10A and F17A (right panel).

To further define critical residues and to assess whether the loop structure is required for antiviral activity, we generated short peptides of six, seven or ten amino acids corresponding to the C- or N-termini of VIR-148, respectively. To determine their antiviral activity, we pretreated TZM-bl reporter cells with various concentrations of individual or mixed peptides and subsequently exposed them to the CXCR4-tropic HIV-1 NL4-3 strain. TZM-bl reporter cells are highly susceptible to HIV-1 infection and contain the β-galactosidase reporter gene under the control of the HIV-1 long terminal repeat.^34^ Three days later, infection was quantified by β-galactosidase assay. Individually, these short peptides did not inhibit HIV-1 infection (Fig. 2A, left). In contrast, mixtures of the split VIR-148 fragments, allowing bridging via the cysteines at their C- and N-termini, inhibited HIV-1 infection almost as efficient as the parental intact VIR-148 peptide. In agreement with these results, ReaxFF-based molecular modelling confirmed the ability of the linked N- and C-terminal VIR-148 fragments to stably bind the HIV-1 gp41 FP (Fig. 2A, right). Mass Spectrometry confirmed the formation of the heterologous disulfide-bridged LEAIPC-CVAFNKPFVF VIRIP derivative from the two short peptides (Fig. 2B). Altogether, these results showed that the N- and C-terminal regions of VIR-148 are both required and sufficient for antiviral activity.

**Figure 2.**
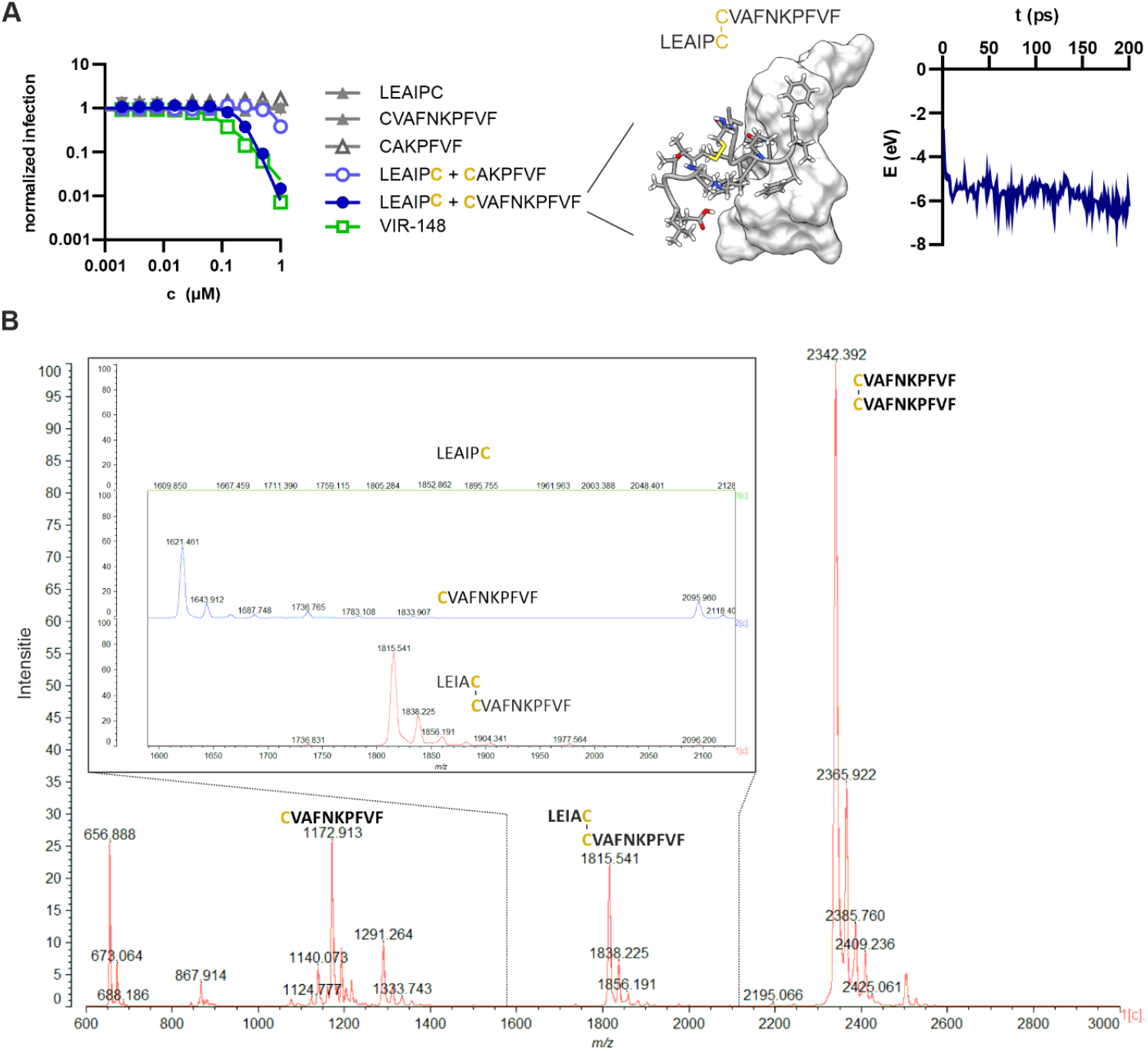
Short N- and C-terminal fragments linked by a cysteine-bridge show efficient anti-HIV-1 activity. (A) TZM-bl reporter cells were incubated with increasing concentrations of VIR-148 or the indicated individual or mixed N- and C-terminal fragments (left). Cells were subsequently infected with HIV-1 NL4-3, and beta-galactosidase activity was determined three days post-infection. ReaxFF-based molecular dynamics simulation of the interaction between the indicated C-C-linked short VIR-148 fragments and the HIV-1 gp41 FP (middle). Trajectory after 150 ps (left) and ReaxFF MD simulation of the cysteine-bridge-coupled split VIR-148 peptides (right). (B) Mass spectrometry analysis of an equimolar mix of the two cysteine-containing VIR-148 fragments in cell culture medium.

After confirming that the loop structure encompassing the internal residues of VIR-148 is not critical for FP binding and antiviral ability, we sequentially deleted the central region of the peptide, thereby gradually reducing its size (Fig. 3A). For comparison, we included VIR-102, the most active VIRIP derivative that does not contain a cysteine and is therefore unable to dimerize or stabilize its structure through a disulfide bridge^26^. The internally truncated VIRIP (itVIR) derivatives itVIR4, 5, and 6 showed high antiviral activity (Fig. 3A). However, these derivatives formed visible precipitates and exhibited hemolytic activity (Fig. 3B). In contrast, derivatives itVIR-7 through itVIR-13 were well-soluble and did not exhibit hemolytic activity (Fig. 3B) while maintaining IC_50_ values close to their VIR-148 precursor (Fig. 3A). itVIR-13 (named “size-optimized” soVIRIP hereafter) had only half the size (10 amino acids) and mass (1.11 kDa) compared to the parental VIR-148 peptide but maintained full antiviral activity (Table 1).

**Table 1.**
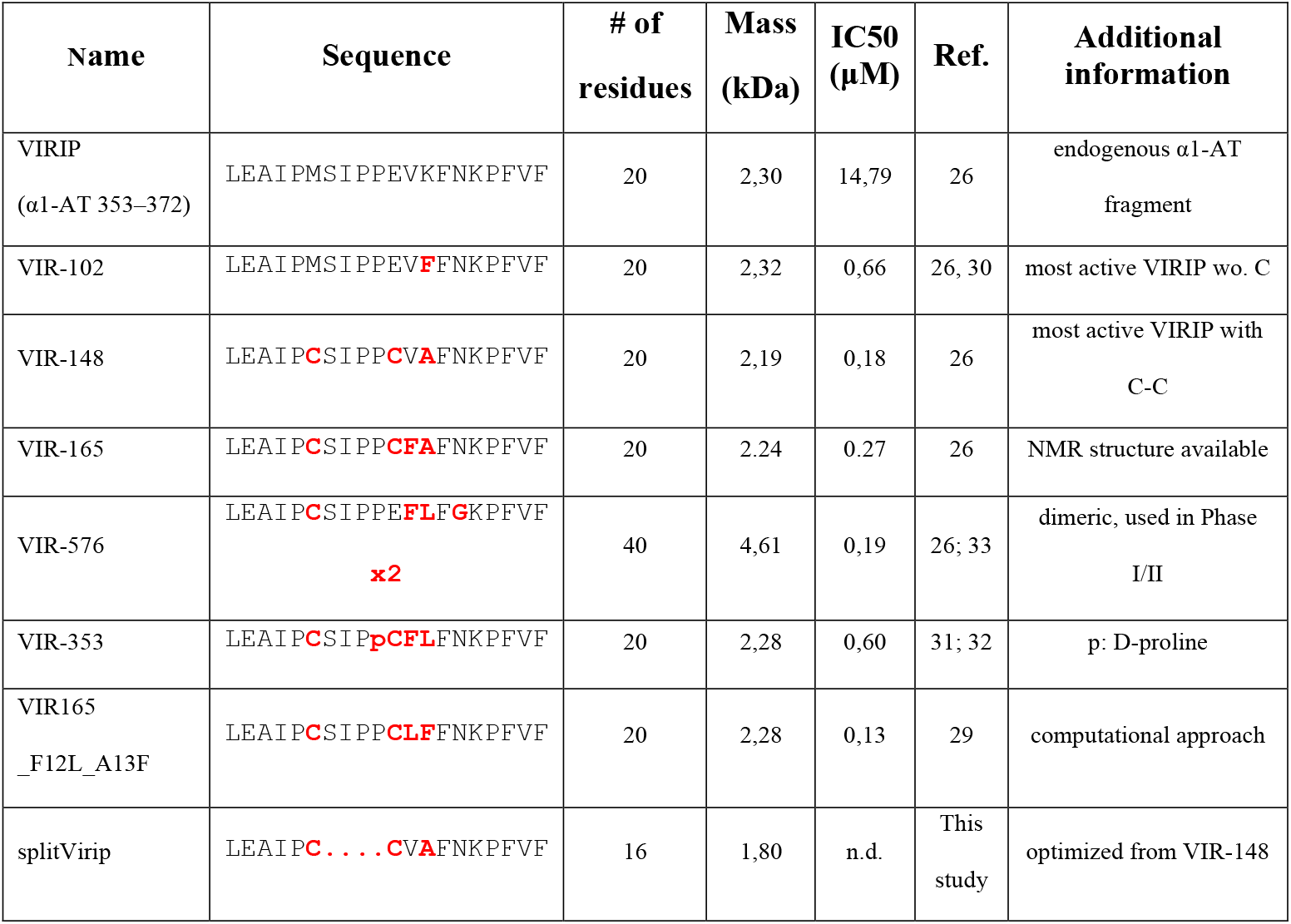
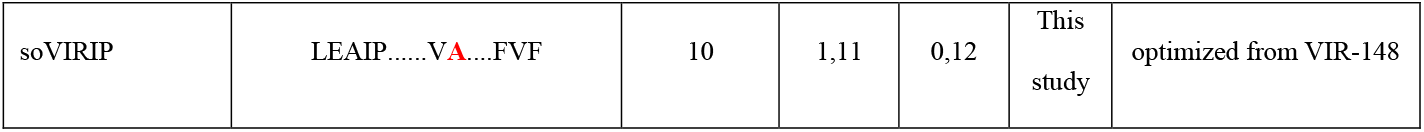
Overview on VIRIP derivatives.

**Figure 3.**
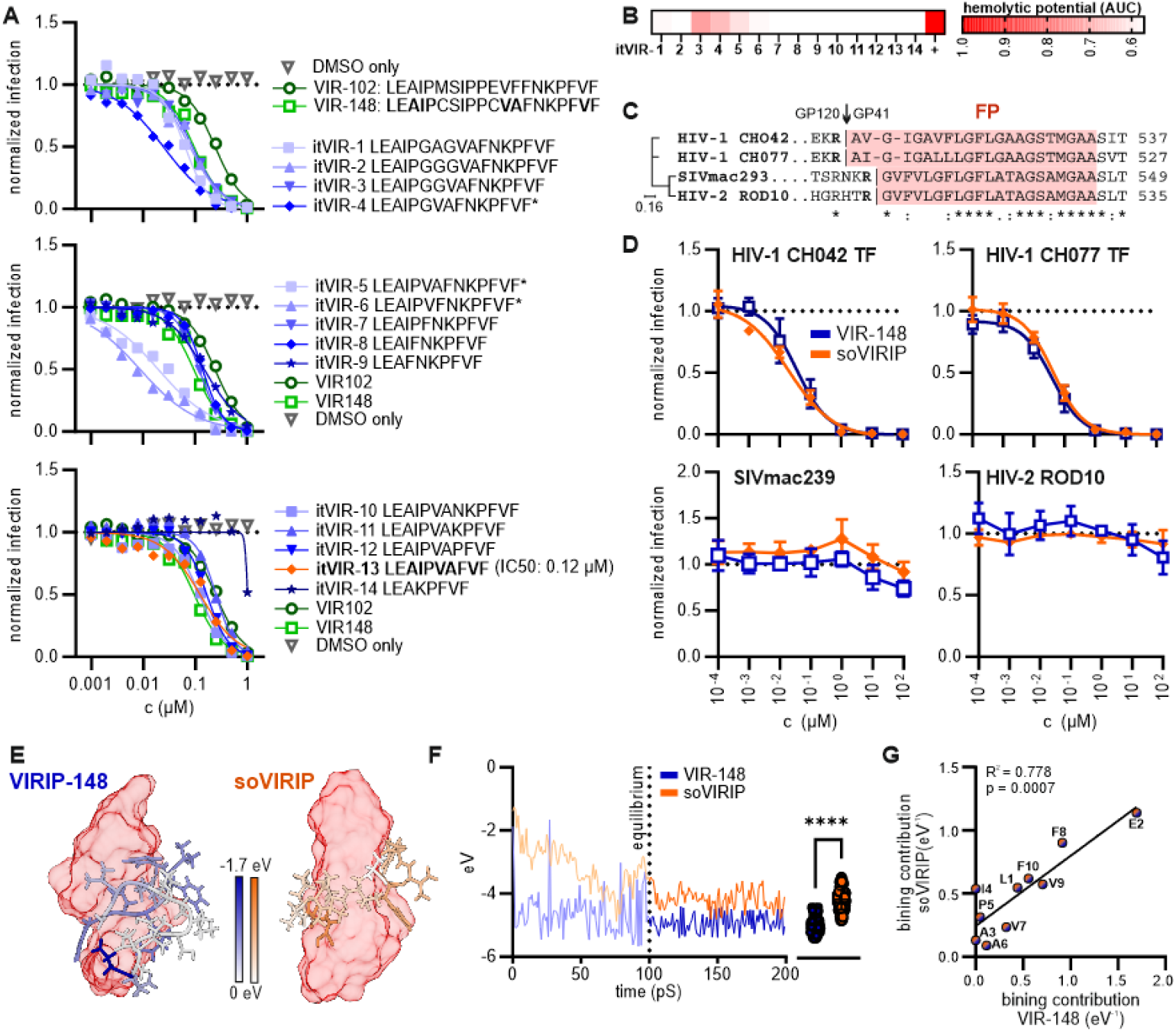
Size optimization of VIR-148. (A) TZM-bl reporter cells were incubated with increasing concentrations of VIR-148, VIR-102, or the indicated internally truncated itVIRIP derivatives. Cells were subsequently infected with HIV-1 NL4-3, and beta-galactosidase activity was determined three days after infection. (B) Hemolytic effect of indicated compounds on human erythrocytes. Each tile represents the area under the curve (AUC) for hemolytic activity. Full haemolysis is represented by a Triton-X control. (C) Amino acid alignment of the FP regions (red) of HIV-1 CH042, CH077, SIVmac239 and HIV-2 ROD10. (D) TZM-bl cells were incubated with increasing concentrations of VIR-148 or soVIRIP and subsequently infected with the HIV-1 CH042 or CH077, SIVmac239 or HIV-2 ROD10. (E) Exemplary ReaxFF trajectory of VIR-148 (blue) and soVIRIP (orange) binding to the HIV-1 gp41 FP (surface, red) after 150 ps of simulation time. The contribution of each amino acid to the binding interaction is indicated by color saturation. (F) Time-resolved ReaxFF-based molecular dynamics simulation of the interaction between soVIRIP and the HIV-1 FP. The trajectories are shown for VIR-148 (blue) and soVIRIP (orange). (G) Correlation between the free energy contribution of each amino acid of soVIRIP with the corresponding amino acid in VIR-148, based on ReaxFF simulations.

Further internal truncations, such as in itVIR-14, strongly impaired antiviral activity (Fig. 3A, bottom). It has been previously shown that VIRIP and its derivatives inhibit diverse HIV-1 strains but are inactive against HIV-2 and simian immunodeficiency viruses infecting macaques (SIVmac),^26^ which show substantial amino acid diversity in the FP region from that of HIV-1 domain (Fig. 3C). In agreement with the previous data, both the original VIR-148, as well as the soVIRIP derivative, efficiently inhibited the clade B CH077 and clade C CH042 transmitted-founder (TF) HIV-1 strains with IC_50_ values of 39 nM and 37 nM, respectively (Fig. 3D). In contrast, both VIRIP derivatives were inactive against HIV-2 ROD10 and SIVmac239, suggesting that the determinants of VIR-148 and soVIRIP interaction with the HIV-1 gp41 FP are conserved. This was confirmed by further ReaxFF-based modelling of VIR-148 and soVIRIP binding to the HIV-1 gp41 FP (Fig. 3E). After an equilibration time of 100 ps, the total free energy of soVIRIP binding to the HIV-1 FP decreased by 0.25 eV (Fig. 3F). Notably, the contribution of individual amino acid residues in VIR-148 and soVIRIP to gp41 FP binding correlates significantly (Fig. 3G). Altogether, our *in silico* analyses and *in vitro* results suggest that VIR-148 and short VIRIP target the same region in the HIV-1 gp41 FP and bind with similar efficiency.

To assess potential toxic effects of VIRIP and its optimized derivatives *in vivo*, we used zebrafish embryos. This model is increasingly used in toxicity studies since most zebrafish organs perform the same functions as their human counterparts. The transparency of the embryos allows for evaluation not only of mortality, but also of sublethal cytotoxicity (necrosis, lysis), developmental toxicity (developmental delay, malformations) or toxicity affecting specific organ systems, in particular cardiotoxicity (heart edema, reduced circulation) and neurotoxicity (reduced touch escape response).^35^ We found that the optimized soVIRIP derivative did not display significant toxic effects in the zebrafish model, whereas NRC-03 (cytotoxic control) and abamectin (neurotoxic control) exhibited the expected toxicities (Fig. 4). Our finding that soVIRIP was non-toxic at concentrations about 2,500-fold higher than the IC_50_ against primary HIV-1 strains suggests that it might be well tolerated *in vivo*.

**Figure 4.**
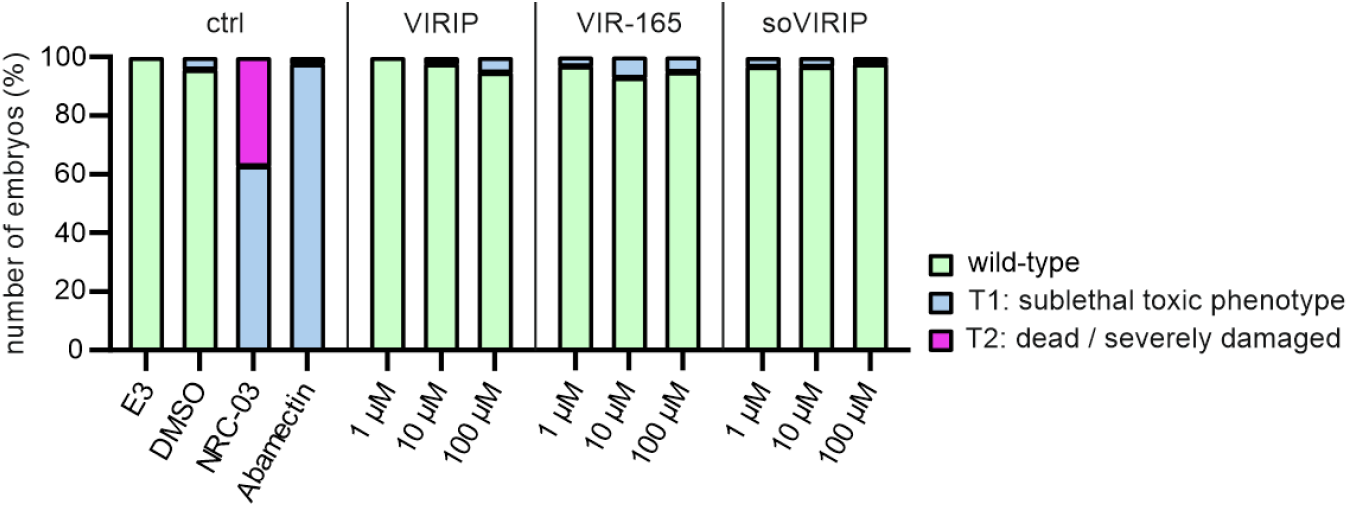
soVIRIP is not toxic to embryonic zebrafish. Twenty-four hours post-fertilization, dechorionated zebrafish embryos were exposed to DMSO, NRC-03 (cytotoxic control), abamectin (neurotoxic control), or increasing amounts of the indicated peptide for 24 hours. Data shown are derived from 60 embryos per group, sampled in two independent experiments.

In the present study, we used ReaxFF-based prediction of amino acids involved in the binding of an optimized VIRIP derivative to the HIV-1 FP to optimize the size of this antiviral peptide fusion. Prediction of the key residues critical for binding by this computational approach allowed us to reduce the size of the antiviral peptide by half without compromising its activity against HIV-1 (Table 1). Such a reduction in peptide size can enhance therapeutic potential by improving its bioavailability, stability, and ease of synthesis, while maintaining the crucial interactions required for antiviral efficacy. Notably, membrane permeability and hence oral availability is size dependent and rapidly increases for peptides with molecular mass of ≤1 kDa,^36^ which is approximated by soVIRIP (1.1 kDa). Our results provide proof-of-concept that ReaxFF-based predictions of target binding are a powerful tool for the optimization of peptides for clinical applications. Overall, this approach may allow to reduce the production costs and immunogenicity of other therapeutic peptides, while retaining full activity.

## Material and Methods

### Cell culture

HEK293T cells were provided and authenticated by the ATCC. TZM-bl reporter cells were provided and authenticated by the NIH AIDS Reagent Program, Division of AIDS, NIAID. HEK293T and TZM-bl cells were maintained in Dulbecco’s modified Eagle medium (DMEM) supplemented with FCS (10%), L-glutamine (2 mM), streptomycin (100 mg/mL) and penicillin (100 U/mL). Cells were cultured at 37°C, 90% humidity and 5% CO2.

### Virus stock production

Virus stocks were generated by transient transfection of HEK293T cells using the calcium-phosphate precipitation method. One day before transfection, 0.8 × 10^6 HEK293T cells were seeded in 6-well plates (Sarstedt, Germany). At a confluence of 80% cells were used for transfection. For the calcium-phosphate precipitation method, 5 μg of the proviral HIV-1 constructs NL4-3,^37^ CH055_TF1, CH077_TF1,^38^ as well as HIV-2 ROD10^39^ and SIVmac239^40^ were mixed with 13 μl 2 M CaCl2 and the total volume was filled up to 100 μl with water. This solution was added dropwise to 100 μl of 2x HBS. The transfection cocktail was vortexed for 5 sec and added dropwise to the cells. The transfected cells were incubated for 8-16 h before the medium was replaced with fresh supplemented DMEM. 48 h post transfection, virus stocks were prepared by collecting the supernatant and centrifuging it at 1300 rpm for 3 min.

### TZM-bl infection assays

To determine infectious virus yield, 10,000 TZM-bl reporter cells/well were seeded in 96-well plates and infected with cell culture supernatants, under reduced 1% FCS conditions, in triplicates on the following day. Three days p.i., cells were lysed and β-galactosidase reporter gene expression was determined using the GalScreen Kit (Applied Bioscience) according to the manufacturer’s instructions with an Orion microplate luminometer (Berthold).

### Peptides

All peptides were commercially obtained from KE Biochem (China) with a purity of ≥95%. The peptides were dissolved in dimethyl sulfoxide (DMSO, Sigma-Aldrich, Hamburg, Germany) at a stock concentration of 50 mM and further diluted in phosphate-buffered saline (PBS) or Dulbecco’s Modified Eagle Medium (DMEM) cell culture media before use.

### Infection Inhibition

10,000 TZM-bl reporter cells were seeded in 96-well plates in 100 μl of supplemented DMEM. The following day, the medium was changed to reduced 1% FCS-supplemented DMEM, and serially diluted peptide was added to the cells. The cells were subsequently infected with 10 μl of infectivity-normalized virus stocks. At 3 days post-infection, the supernatant was removed, and β-galactosidase activity was measured as previously described.

### ReaxFF based molecular dynamics simulations

The molecular dynamics simulations of this study use the ReaxFF approach,^41^ which includes bond order-dependent energy terms that dynamically adapt to the local atomic environment. The C/H/O/N glycine/water parameters developed by Rahaman *et al*. and extended by Monti *et al*. were used.^42,43^ Rigorous tests confirmed the accuracy and transferability of the force field with a training set based on DFT-B3LYP/6-311++G** calculations of amino acid structures.^42,44^ The simulations were also compared against the classical ff99SB force field from the AMBER family.^45^ Based on the structure of the complex between the HIV-1 gp41 FP and VIR-165 from the Protein Data Bank^46^: 2JNR (https://www.rcsb.org/structure/2JNR), the initial atomic positions were obtained.

Equilibration (300 K for 0.5 ns) was performed by ReaxFF simulations (reactive molecular dynamics) within the Amsterdam Modelling Suite 2020 (http://www.scm.com). Based on the equilibrated structure, the amino acids from VIRIP were replaced by corresponding amino acids. After additional equilibration (300 K for 0.5 ns), ReaxFF (reactive molecular dynamics) simulations were performed within the NVT ensemble over 25 ps while the system was coupled to a Berendsen heat bath (T = 300 K with a coupling constant of 100 fs). The interaction energy was obtained by averaging over these simulations. UCSF Chimera^47^ was used for all visualisations.

### Hemolysis assay

Fresh human blood, collected by venipuncture, was centrifuged (10 min, 1000 xg, 4 °C) to pellet erythrocytes, which were then washed three times and resuspended (1:10) in DPBS. Peptides were serially diluted in a 96-well plate in DPBS. To 90 μl of peptide dilution, 10 μl of erythrocyte suspension was added. Following a 1-h incubation at 37 °C, while shaking at 500 rpm, the plates were centrifuged (5 min, 1000 xg, 4 °C). Supernatant was transferred to transparent 96-well plates for absorbance measurement at λ = 405 nm, with full erythrocyte lysis indicated by the Triton-X control.

### MSA

Alignments of viral FP amino acid sequences were performed in Clustal Omega (https://www.ebi.ac.uk/Tools/msa/clustalo/) using the ClustalW 63 algorithm and an ordered input. The resulting phylogenetic tree was transferred to ITOL (https://itol.embl.de/) and visualized as a rectangular phylogenetic tree in default settings.

### Mass spectrometry

Single or equimolarly mixed amounts of peptide, diluted in serum-reduced DMEM, were spotted on a 384-well plate and analyzed with an Axima Confidence MALDI-TOF mass spectrometer (Shimadzu, Japan) as follows: plate wells were coated with one microliter of 10 mg/ml CHCA (matrix) in HFBA/acetonitrile/2-propanol/water (v/v: 2.5/25/25/47.5), and the solvent evaporated; then, the sample (0.5 µL) was mixed with matrix (0.5 µL) onto the dry pre-coated well, and the solvent evaporated. The measurement was done in linear mode. An accelerating voltage of 20 kV was applied to the ion source, and the laser shots were automatically done following a regular circular raster of a diameter of 2,000 µm and spacing of 200 µm on every well; 100 profiles were acquired per sample, and 20 shots were accumulated per profile. The measurement and MS data processing were performed with MALDI-MS Application Shimadzu Biotech Launchpad 2.9.8.1 software (Shimadzu, Japan).

### Zebrafish

For *in vivo* studies, wild-type zebrafish embryos (Danio rerio) were dechorionated at 24 hours post fertilization (hpf) using digestion with 1 mg/ml pronase (Sigma) in E3 medium (83 μM NaCl, 2.8 μM KCl, 5.5 μM 202 CaCl2, 5.5 μM MgSO4). In a 96-well plate, 3 embryos per well were exposed for 24 h to 100 µL of E3 containing VIRIP peptides at the concentrations indicated in the figures. Two independent assays were performed, each with 10 × 3 embryos. The peptide solvent (DMSO), diluted in E3, was used as negative control at the same amount as introduced by the peptide stock. As positive control for acute toxicity/cytotoxicity the pleurocidin antimicrobial peptide NRC-03 (GRRKRKWLRRIGKGVKIIGG AALDHL-NH2) was used at a concentration of 3 µM as described.^48^ Abamectin at a concentration of 3.125 µM was used as positive control for neurotoxicity.^49^ At 48 hpf (after 24 h of incubation) embryos were scored in a stereomicroscope for signs of cytotoxicity (lysis and/or necrosis), developmental toxicity (delay and/or malformations) or cardiotoxicity (heart edema and/or reduced or absent circulation). Each embryo was also touched with a needle and reduced or absent touch response (escape movements) was evaluated as signs of neurotoxicity if and only if no signs of acute toxicity were present in the same embryo. Embryos were categorized within each of these toxicity categories into several classes of severity.^35^ Chi-Square test was used to calculate whether the distribution of embryos into toxicity classes differed significantly between the PBS negative control and the test substances.

## Corresponding Author

Fabian Zech

## Author Contributions

Conceptualization, F.Z. and F.K; funding acquisition, F.K.; investigation, F.Z., C.J., A.R.A., J.K., L.S., G.W. and T.J.; writing original draft, F.Z. and F.K.; reviewing, editing and approving the final version: all authors.

## ACKNOWLEDGMENT

The authors acknowledge support by the state of Baden-Württemberg through bwHPC and the German Research Foundation (DFG) through grant no INST 40/575-1 FUGG (JUSTUS 2 cluster).

## ABBREVIATIONS

ACE2: angiotensin-converting enzyme 2
AUC: area under the curve
CD4: cluster of differentiation 4
CXCR4: C-X-C chemokine receptor type 4
DMEM: Dulbecco’s modified Eagle medium
DMSO: dimethyl sulfoxide
DNA: deoxyribonucleic acid
eV: electron volt
FP: fusion peptide
FCS: fetal calf serum
GP41: glycoprotein 41
hpf: hours post fertilization
HIV: human immunodeficiency virus
HIV-1: human immunodeficiency virus type 1
HIV-2: human immunodeficiency virus type 2
IC50: half maximal inhibitory concentration
kDa: kilodalton
MS: mass spectrometry
NMR: nuclear magnetic resonance
PBS: phosphate-buffered saline
ps: picosecond
ReaxFF: reactive force field
SARS-CoV-2: severe acute respiratory syndrome coronavirus 2
SAR: structure-activity relationship
SIVmac: simian immunodeficiency virus infecting macaques
soVIRIP: size-optimized Virus Inhibitory Peptide
T20: Enfuvirtide (Fuzeon HIV-1 entry inhibitor)
TF: transmitted-founder (virus strain)
TZM-bl: reporter cell line containing β-galactosidase under the HIV-1 LTR control
UCSF: University of California San Francisco
VIRIP: Virus Inhibitory Peptide

